# BTK and MMP9 regulate NLRP3 inflammasome-dependent cytokine and NET responses in primary neutrophils

**DOI:** 10.1101/2024.02.23.581733

**Authors:** Vinicius N. C. Leal, Francesca Bork, Juli-Christin von Guilleaume, Carsten L. Greve, Stefanie Bugl, Bettina Danker, Zsofía A. Bittner, Bodo Grimbacher, Alessandra Pontillo, Alexander N. R. Weber

## Abstract

**Background:** Inflammation is a double-edged state of immune activation that is required to resolve threats harmful to the host but can also cause severe collateral damage. Polymorphonuclear neutrophils (PMN,) the primary leukocyte population in humans, mediate inflammation through the release of cytokines and neutrophil extracellular traps (NETs). Whilst the pathophysiological importance of NETs is unequivocal, the multiple molecular pathways driving NET release are not fully defined. Recently, NET release was linked to the NLRP3 inflammasome which is regulated by Bruton’s tyrosine kinase (BTK) in macrophages.

**Objective:** As NLRP3 inflammasome regulation by BTK has not been studied in neutrophils, we here explored a potential regulatory role of BTK in primary murine and human neutrophils and matched monocytes or macrophages from Btk-deficient vs WT mice or healthy donors (HD) vs *BTK* deficient X-linked agammaglobulinemia (XLA) patients, respectively.

**Methods:** Cytokine, MPO and MMP-9 release were quantified by ELISA, NET release and inflammasome formation by immunofluorescence.

**Results:** Surprisingly, in both mouse and human primary neutrophils, we observed a significant increase in NLRP3 inflammasome-dependent IL-1β and NETs when BTK was absent or inhibited, whereas IL-1β release was decreased in corresponding primary mouse macrophages or human PBMC, respectively. This suggests a negative regulatory role of BTK in neutrophil NLRP3 activation. Both IL-1β and NET release in mouse and human primary neutrophils were strictly dependent on NLRP3, caspase-1 and, surprisingly, MMP-9.

**Conclusion:** This highlights BTK and MMP-9 as novel and versatile inflammasome regulators and may have implications for the clinical use of BTK inhibitors.

**Key messages:**

- Neutrophils contribute to inflammation by release of interleukin-1β and Neutrophil Extracellular Traps (NETs) via the NLRP3 inflammasome
- Bruton’s tyrosine kinase (BTK) is a negative regulator of NLRP3-mediated primary human neutrophil functions, whereas it positively regulates NLRP3 in monocytes
- MMP-9 is both effector and regulator of the neutrophil NLRP3 inflammasome

**Capsule summary:** Here we report that interleukin-1β and Neutrophil Extracellular Traps (NETs) release via the NLRP3 inflammasome is negatively regulated by Bruton’s tyrosine kinase (BTK) in primary neutrophils. Thus, targeting BTK using FDA-approved inhibitors might increase neutrophil functions.

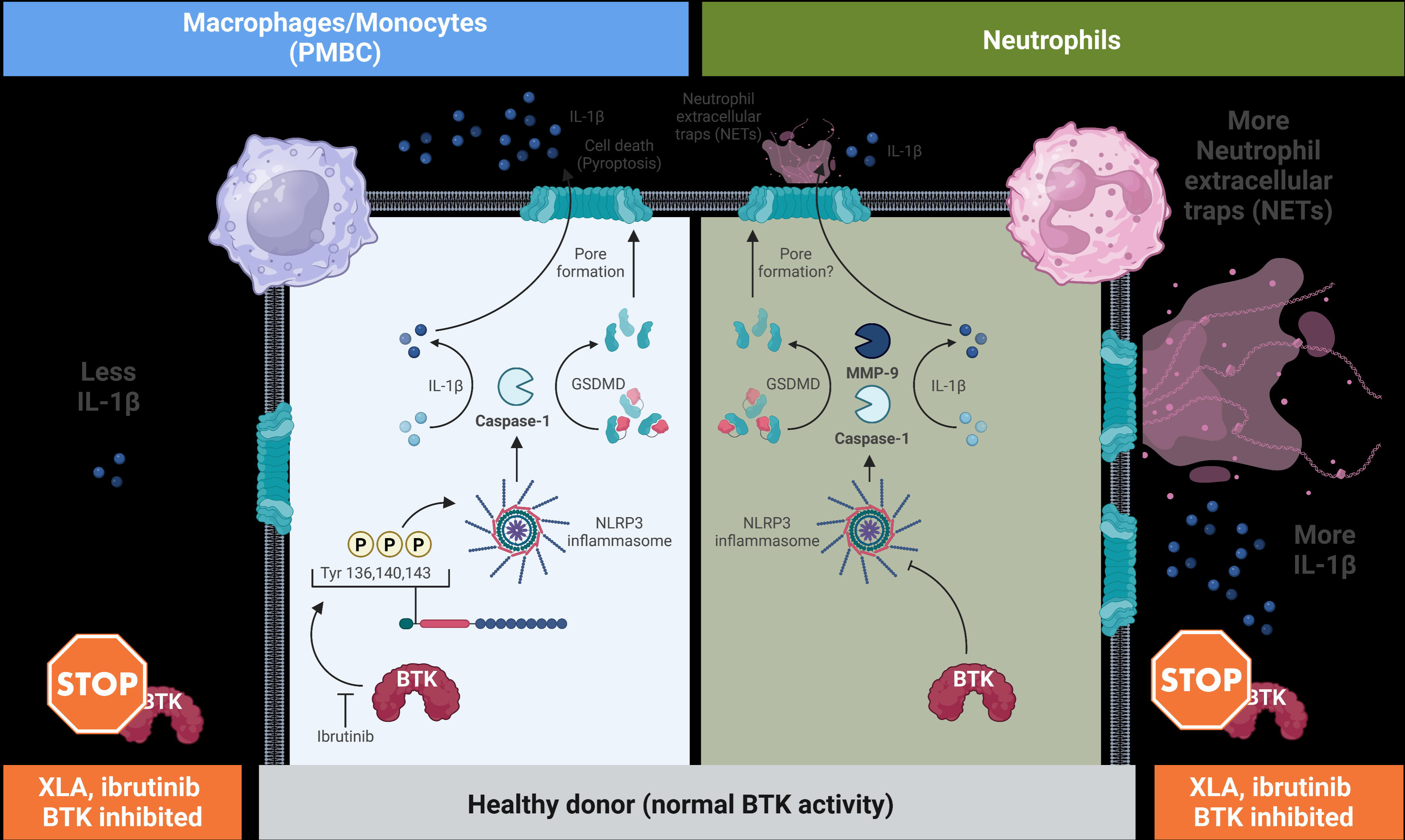

## Introduction

Inflammation is a state of immune activation associated with most of the top ten leading causes of death globally, including ischemic heart disease, stroke, lung disease, neurodegeneration and infections^1^. Inflammation is typically the result of an acute threat, e.g. an infection or tissue damage, and an attempt to increase local immune activity and perfusion to quickly resolve the threat. Thus, inflammation is typically well-constrained and transient. However, even transient inflammation, e.g. in heart or brain ischemia, may have adverse long-term effects, or inflammation may turn chronic. Inflammation is typically initiated or maintained via soluble inflammatory mediators released by immune cells such as leukocytes in response to so-called microbe-associated molecular patterns (MAMPs) or sterile danger-associated molecular patterns (DAMPs). MAMPs and DAMPs act via so-called pattern recognition receptors (PRRs), cellular danger sensors of the innate immune system^2^. PRR signaling drives the production and/or release of soluble inflammatory mediators such as the cytokines Interleukin (IL)-1β, IL-6 or TNF or prostaglandins from macrophages or polymorphonuclear neutrophilic granulocytes (PMN). The latter can unleash pre-stored inflammatory mediators from pre-stored granula, but also release so-called neutrophil extracellular traps (NETs)^3,4^. NETs are an ancient antimicrobial mechanism aimed at trapping and killing microorganisms^5^, but they also contain potent DAMPs such as the newly identified NET-associated naRNA (^6^ and pre-print ^7^). Given their large abundance in the blood (∼65%), PMN are a potentially pro-inflammatory cell population to be reckoned with and hence have been implicated in debilitating or even life-threatening inflammatory conditions such as psoriasis, rheumatoid arthritis and COVID-19 ^6, 8, 9^. Understanding molecular pathways governing the release of inflammatory mediators or NETs from neutrophils is therefore critical for both our understanding of inflammatory conditions as well as their treatment.

Both IL-1β and NET release in human and murine neutrophils were recently found to be regulated by the NLRP3 inflammasome ^4^, a multi-protein complex best studied in macrophages (reviewed in Ref. 10). In macrophages the NLRP3 inflammasome consists of the sensor, NLRP3, an adaptor, ASC, and an executing enzyme, caspase-1. Caspase-1, is activated by proximity-induced autoactivation within inflammasome complexes and then processes substrates like pro-inflammatory cytokines (e.g. IL-1β and IL-18) or Gasdermin D, which induces a type of cell death termed pyroptosis^11^. In macrophages, nigericin, a K^+^/H^+^ ionophore antibiotic derived from *Streptomyces hygroscopicus*, extracellular ATP or pore-forming toxins ^12, 13^, like Staphylococcal LukAB ^14^, directly or indirectly affect K^+^ efflux, a signal that leads to a conformational change in NLRP3 ^15^ and the assembly of inflammasomes ^16, 17^. These can form via two pathways, a rapid pathway kickstarted by pre-formed oligomeric ‘cages’ of NLRP3 at the trans-Golgi network (TGN) and microtubule organizing center (MTOC) ^16^; and a recently described slower, membrane-independent pathway relying on non-cage NLRP3 (pre-print ^18^). In the TGN/MTOC pathway, activation of NLRP3 requires membrane association and dissociation, processes that are regulated by certain kinases, including Bruton’s tyrosine kinase (BTK) ^14, 19, 20^. BTK is well known for its function in B cells, as a target for B cell malignancies, and as the genetic cause of X-linked agammaglobulinemia (XLA), an immunodeficiency characterized by the almost complete lack of mature B cells and hence antibodies ^21^. The XLA phenotype is recapitulated in *Btk* KO and so-called *Xid* mice. *Xid* comprises a *Btk* C to T substitution at coding nucleotide 82, changing codon 28 from an arginine to a cysteine (p.R28C) in the kinases’s pleckstrin homology (PH) domain. R28C prevents membrane-association, a requirement for BTK activation ^22^. In addition to its role in B cells, in macrophages BTK was shown to promote NLRP3 inflammasome activity, enabling maximal IL-1β release in murine and human cells. Consequently, XLA patients and individuals receiving FDA-approved BTK inhibitors, e.g. ibrutinib, showed reduced IL-1β release ex vivo ^14^. A positive regulatory role of BTK was also observed in in experimental murine models of NLRP3 activation, e.g. stroke ^23^, acute lung Injury ^24^, sepsis ^25^ and neuroinflammation ^26^. On the molecular level, BTK interacts constitutively with NLRP3 in macrophages and promoted TGN dissociation and NLRP3 oligomerization via direct phosphorylation of NLRP3 at three adjacent tyrosine residues ^19^. Thus, at physiologically low LPS concentrations, BTK boosts NLRP3 activity ^4, 14, 23^, whereas at high LPS concentrations, BTK was also found to suppress NLRP3 activity ^27^. We previously proposed BTK may thus be a versatile rheostat able to regulate NLRP3 depending on further signals experienced by macrophages ^28^.

In neutrophils the main players and the mechanism of NLRP3 activation and specifically the role of BTK have not been investigated systematically. However, Karmakar et al., ^29^ showed that both, human and murine neutrophils have a functional NLRP3 inflammasome, as evidenced by IL-1β release after LPS priming and ATP stimulation ^29^. Similar to what occurs in macrophages, the pore forming toxin nigericin was also able to induce IL-1β release in LPS primed neutrophils ^30, 31^. Furthermore, previous reports showed NLRP3 activation in infections with *Staphylococcus aureus* ^32^, *Streptococcus pneumoniae* ^33^ and SARS-CoV-2 ^34, 35^. Moreover, previous studies linked NET release to NLRP3 activation in neutrophils exposed to nigericin and naRNA-LL37 complexes ^4, 30, 34, 35^. However, how exactly NLRP3-mediated NET release occurs and is regulated remains elusive. The role of GSDMD as a possible player in NET release has been intensely discussed already ^30, 31, 35^, but BTK has not been investigated even though it is highly expressed in neutrophils ^36^.

Apart from NET release, NLRP3 activation in neutrophils has been linked to the release of azurophilic/primary and gelatinase/tertiary granules ^37^. The latter contain the matrix metalloproteinase 9 (MMP-9; also known as gelatinase B) and a connection between MMP-9 release and NLRP3 activation was previously shown in both, mouse ^38-40^ and human cells ^35^. On the other hand, MMP-9 could potentially act in IL-1β processing ^41, 42^. Hence, a role for MMP-9 not only as effector but also as a regulator/executioner of NLRP3 activation pathway in neutrophils is thus conceivable.

Since BTK and MMP-9 inhibitors are in clinical use or development ^43, 44^ and neutrophils are an essential and abundant white blood cell population with inflammasome capacity, we sought to investigate the role of BTK in primary neutrophils. We found that in primary murine and human neutrophils genetic BTK ablation or BTK inhibitors increased NLRP3 activity in terms of IL-1β release output, conversely to matched bone marrow-derived macrophages (BMDM) or peripheral blood mononuclear cells (PBMC) from *Btk*-deficient or X-linked agammaglobulinemia (XLA) patients with verified clinical *BTK* deficiency, respectively. Importantly and in line with IL-1β release, NET formation was increased when BTK activity was suppressed. This suggests a novel inhibitory role of this kinase in the regulation of neutrophil NLRP3 inflammasome. Interestingly, both IL-1β and NET release in mouse and human primary neutrophils were strictly dependent not only on NLRP3, caspase-1 but also, surprisingly, MMP-9. This highlights BTK and MMP-9 as novel and versatile NLRP3 inflammasome regulators in primary neutrophils.

## Methods

### Reagents

PMA (tlrl-pma), nigericin (tlrl-nig), LL37 (tlrl-l37), as well as the PRR ligands LPS (tlrl-peklps), R848 (tlrl-r848), TL8 (tlrl-tl8506), MDP (tlrl-mdp), Pam2CSK4 (tlrl-pm2s-1) and the TLR8-inhibitor CU-CPT9a (inh-cc9a) were from Invivogen, Ionomycin was acquired from Sigma (I0634-1MG). RNase inhibitor (N2615) was from Promega, RNase A (EN0531), DNase I (EN0521) and DNase inhibitor (EN0521) were from Thermo Fisher. The PAD4-inhibitor Cl-amidine (506282) was from Merck Millipore. The NLRP3-inhibitor MCC950 was from Cayman chemicals (Cay17510-1) and the GSDMD-inhibitor disulfiram from Medchem Express (HYB0240). Further reagent and kits are listed in Supplementary Table S1, buffers in Supplementary Table S2.

### Human subjects

XLA patients (n = 4) and matched HD (n = 4) were recruited at the Centre of Chronic Immunodeficiency at the University Hospital Freiburg. XLA clinical and genetic identification was performed as described in ^14^. For additional experiments with primary human cells, 20 healthy blood donors were recruited at the Interfaculty Institute of Cell Biology, Department of Immunology, University of Tübingen (Germany) and at the “Instituto de Ciências Biomédicas (ICB) of Universidade de São Paulo (USP)” in Brazil. In each case, approval for use of biological material was obtained by the local ethics committee of the Medical Faculty Tübingen and ICB/USP, and written informed consent obtained before blood drawing in accordance with the Declaration of Helsinki and local legislations.

### Animals

Mice maintaining the *Xid* allele on a BL6 background (B6J.CBA-*Btk^Xid^*) were generated by backcrossing of the original CBA/CaHN-*Btk^Xid^*/J (Jackson stock no. 001011) mice to C57BL/6J for seven generations using speed congenics (GVG genetic monitoring, Leipzig, German) until autosomal and sex chromosomes were >98% congenic with the C57BL/6J strain whilst monitoring the *Xid* allele by genotype-specific PCR. Subsequently, these *Xid* mice were maintained locally in specific pathogen–free conditions. The same applied to *Btk* KO (B6;129S-*Btk^tm1Wk^*, originally from Khan et al., 1995 ^45^), *lrp3* KO (B6.129S6-*Nlrp3^tm1Bhk^*/J, Jackson stock no. 021302Laboratory), and WT C57BL/6J (Jackson) mice. All animals used were from both sexes and with the age ranging from 8 to 14 weeks. All animal experiments were approved by local authorities and performed in accordance with local institutional guidelines.

### Primary human neutrophil and PBMC isolation

EDTA-blood samples were diluted 1:2 in PBS and then layered on top of Ficoll (1.077 g/mL) and centrifuged for 25 min at 509 x g at Room Temperature (RT) without brake and acceleration. Afterwards, the PBMCs were isolated and then everything but the red pellet was discarded, and the pellet resuspended carefully in 1x erythrocyte lysis buffer (Ammonium-Chloride-Potassium/ACK), filling up to 50 ml, to lyse the red blood cells (RBC). The lysis was performed for 20 min on a rolling shaker at 4 °C. Another centrifugation step at 300 x *g* for 10 min (no brake, no acceleration) was performed and the supernatant discarded. A second RBC lysis was performed with 25 ml ACK, on the roller shaker for 5 min at 4 °C. After a final centrifugation step with the same settings, the PMN pellet was resuspended in neutrophil culture medium and 1x10^6^ seeded in 24 well plates containing Poly-L-Lysin coated coverslips for immunofluorescence or 8.25 x 10^5^ seeded in 48 well plates for other assays. For PBMCs, 5 x 10^5^ were seeded in 24 well plates. All cells were incubated at 37 °C and 5% CO_2_ for 30 min before further stimulations. The list of reagents and buffers used is present in Supplementary Tables S1, S2.

### Isolation of primary bone-marrow-derived polymorphonuclear neutrophils (BM-PMNs)

BM-PMNs were isolated from the bone marrow as described^19^. In summary, bone marrow was flushed out from the bones with BM medium, and the neutrophils were isolated using magnetic separation (mouse neutrophil isolation kit, Miltenyi Biotec) following the manufacturer’s instructions. In total, 1 × 10^6^ cells/well were seeded in 24 well plates and maintained resting overnight at 37 °C and 5% CO_2_, prior to stimulation.

### Bone-marrow-derived macrophages (BMDM) differentiation

For BMDM differentiation around 10 x 10^6^ BM cells were resuspended BMDM medium containing 10% granulocyte macrophage colony-stimulating factor (GM-CSF) supernatant and seeded in 10 cm dish plates in a volume of 10 mL RPMI containing 10 % fetal calf serum (FCS), 1% L-glutamine (Gibco) and 1 % Pen Strep (Gibco). After four days of incubation at 37 °C, 5% CO_2_, additional 5 mL of fresh BMDM medium containing GM-CSF were added to the plate and maintained for additional two days.

Finally, the fully differentiated BMDMs were detached with 1 mL phosphate buffered saline (PBS) containing 1 mM ethylenediaminetetraacetic acid (EDTA) and 2% FCS incubated for 15 min. After counting, the cells were resuspended in GM-CSF free BMDM media for further stimulations.

### FACS determination of neutrophil purity and viability

To ensure a highly pure and viable neutrophil preparation, the cells were analysed by FACS after every isolation, checking for specific human and mouse neutrophil markers as well as Live/Dead staining. Briefly, 0.1-1 x 10^6^ cells per well were added in a 96-well U bottom plates and centrifuged for 5 min at 250 x *g*. The staining was performed in 50 μL of FACS buffer containing the appropriate antibodies in a pre-titrated dilution together with Live/Dead (Supplementary Table S1), for 20 min at 4 °C in the dark. Afterwards, cells were washed with 100 μL of FACS buffer and centrifuged again. This was repeated once again, and finally the cells resuspended in FACS buffer and measured right away using a BD FACS Canto II. For later measurements or before intracellular staining, the cells were fixed with 100 μL of Fixation Buffer 1X (Biolegend) for 10 min at RT, then washed and resuspended in FACS Buffer.

For intracellular IL-1β staining, after fixation, the cells were permeabilized with 200 µL of Intracellular Staining Permeabilization Wash Buffer 1X for 5 min, centrifuged 300xg for 5 min and then incubated with the primary antibody for 30 min at 4°C. Then, washed twice with Perm wash solution and analysed in FACS Canto II. Corresponding isotype controls were included for all markers (Supplementary Table S3).

The cells were used when the viability was higher than 85% and the purity was higher than 95% (CD66b^+^CD11b^+^CD15^+^ for human and CD11b^+^Ly6G^+^ for mouse), combined with less than 1% of monocyte contamination (CD14^+^).

### Stimulation and Treatments of PMNs

After resting for 30 min at 37 °C and 5% CO_2_, cells were primed with 10 ng/mL LPS for 3 h for human neutrophils or PBMCs or with 100 ng/mL for 3 h for murine BM PMN or BMDMs. We then pre-treated the cells with different concentrations of selective inhibitors for Btk (Ibrutinib, CGI1746, Acalabrutinib; Selleckchem), Syk (R406; Selleckchem), MMP-9 (JNJ0966; MedChemExpress), Caspase-1 (YVAD; Adipogen) or NLRP3 (MCC950; Cayman) for 30 min, and then stimulated with 5 μM Nigericin (Invivogen) or 5 mM ATP (Sigma) for 1 or 2 h, or with 5 μM LukAB for 1 h ^14^. At the end of the stimulation, the supernatant was collected for ELISA, lactate dehydrogenase (LDH) release and Picogreen NET assays ^46^, and/or the cells fixed and stained for IF. For LDH assay the medium was replaced with OptiMEM (Thermofisher) without FCS prior to the second stimulation.

### ELISA

For cytokine measurements in supernatant or plasma samples, triplicate enzyme-linked immunosorbent assays (ELISAs) were performed using kits provided by BioLegend, R&D or BD Systems (Supplementary Table S1). For human samples, IL-1β, TNF-α, IL-8, IL-6, myeloperoxidase (MPO) and matrix metalloproteinase 9 (MMP-9) were quantified. For murine samples, IL-1β and TNF-α. ELISAs were performed according to the respective manufacturer’s instructions but using high-binding half-area 96-well plates (Greiner). OD measurements were performed using the Molecular Devices Spectramax plus 384 plate reader. The final analysis was done using Microsoft Excel and GraphPad Prism 8.2.0.

### LDH release assay

As a measure of cell death, the amount of LDH released during the *in vitro* experiments were quantified using the kit Cytotoxicity Detection Kit (LDH) provided by Roche. Triton X-100 (Applichem) was used in a final concentration 1%, as a 100% cell death calibrator.

### Picogreen Assay

For NET quantification in neutrophil supernatants or plasma samples, the Quant-iT™ PicoGreen was used in a protocol adapted from ^47^. Briefly, 100 μL/well anti-human MPO antibody (ThermoFisher) pre-diluted 1:500 in PBS was added in a black 96 well plate (Corning, 3603). The plate was then incubated overnight at 4 °C. The next day, the plate was washed 3x with 200 μL washing solution (PBS containing 0.05% saponin) and subsequently blocked with 100 μL blocking solution (PBS+2% BSA) for 2 h at RT. The washing step was then repeated and 100 μL sample per well was added and incubated at 4°C for 4 h or overnight. After another washing step, 100 μL of DNA standard (provided by the kit) and 100 μL of DNA detection reagent was added to all wells. After 5 min of incubation at RT in the dark, the reaction was quantified at 488 nm excitation and 525 nm emission using a BioTek Synergy Neo2 plate reader. The final analysis was done using Microsoft Excel and GraphPad Prism 8.2.0.

### Immunofluorescence Microscopy

After stimulation, the medium was removed and the coverslips rinsed with 500 μL PBS. Then 250 μL of fixation buffer (Biolegend) was added and incubated for 10 min in the dark at RT. The fixation buffer was removed, and the wells rinsed with 400 μL PBS containing 0.05% saponin. Then, 300 μL blocking buffer (PBS containing 10% chicken serum and 0.1% saponin) were added and incubated overnight at 4°C. The primary antibodies (see Supplementary Table S3 for details) targeting H3Cit (1:200), ASC (1:100), MMP-9 (1:200) or Btk (1:200) were diluted in blocking buffer and incubated for 2 h at RT. The wells were then washed 3 times with IF washing solution, and incubated with the respective secondary antibodies (see Supplementary Table S3, diluted 1:500) for 1 h in the dark at RT. Then three washes were performed and 500 µL of Hoechst 33342 (Thermo Fisher; 2 μg/mL) were added for 5 min. Then three further washes were performed, and the coverslips mounted onto microscopy slides using 4 μL of ProLong Diamond Antifade mounting solution. For image acquisition, a Zeiss LSM800 AiryScan confocal fluorescence microscope was used. For NET quantification, images were acquired in Airy Scan mode at 1.5x zoom and frame size 512x512 pixels, 40X with Z-stack acquisition. For ASC, MMP-9 and BTK imaging, the images were acquired in Airy Scan mode, in Super Resolution, 63X. The scan speed was maximum, and no averaging was done. Digital gain was 650-750 V. The laser power was set at 0.5-2%. For NET counting, 3 tiles were acquired per condition. All the images were analysed using ImageJ-Win64.

### DNA isolation and genotyping

The DNA isolation from peripheral blood was carried out using the “Salting Out” method (Miller et al., 1998 ^48^). An NLRP3 gain of function functional germline single nucleotide variant (SNV) was selected (rs10754558) ^49^. Genotyping was performed using allele-specific Taqman® assays (Applied Biosystems, Thermo Fisher Scientific) on a QuantStudio 3 Real-time PCR equipment (Applied Biosystems, Thermo Fisher Scientific).

### Statistical analysis

Experimental data were analyzed using Microsoft Excel 2019 and/or GraphPad Prism 8, microscopy data with ImageJ-Win64 or ZenBlue3 software, flow cytometry data with FlowJo V10. Normal distribution for each group was first assessed using a Shapiro-Wilk normality test in order to select either a parametric or non-parametric test depending on the number of groups (Student’s t-test /Mann Whitney or ANOVA / Friedman), and the type of data (paired/unpaired) using the software GraphPad Prism 8. Multiple testing was accounted for using the appropriate post-hoc tests as described in the figure legends. p < 0.05 was considered statistically significant and is denoted by *p<0.05, **p<0.01, ***p<0.001 and ****p<0.0001.

## Results

### BTK negatively regulates NLRP3-mediated IL-1*β* release in primary human neutrophils

Because we suspected BTK might be a rheostat for NLRP3 activity in monocytes ^14, 19^, we sought to extend current analyses to primary human PMN. Therefore, matched PBMC and PMN from healthy donors (HD) and patients with the *BTK* deficiency, XLA, were analyzed for IL-1β release upon priming with physiologically low (10 ng/ml) LPS and nigericin activation. In confirmation of earlier data, HD PBMC released significantly higher IL-1β levels (presumably from the monocyte fraction, ^19^) than XLA PBMC (**Fig. 1A**), whereas TNF release was comparable (**Fig. 1B**). Conversely, matching HD PMN released significantly lower IL-1β than XLA PMN (**Fig. 1C**) with comparable levels of MPO (**Fig. 1D**), an NLRP3-independent soluble factor released during PMN activation ^50^. Increased IL-1β release in BTK-deficient vs HD PMN was also observed in response to other NLRP3 stimuli, such as ATP and LukAB (**Fig. S1A, B**) and indicated a significantly higher responsiveness of PMN in the absence of BTK, i.e. a negative regulatory role. This was consistent with the observations that unstimulated XLA PMN had higher basal IL-1β levels and spontaneous ASC speck formation (**Fig. S1C-E**), a hallmark of inflammasome activation^51^. To confirm this further, we used the BTK inhibitors, ibrutinib, CGI-1746 and acalabrutinib ^52^, which reduced IL-1β release from PBMC (**Fig. S1H**) as previously described ^14, 19^. Interestingly, concentrations as low as 1 nM, which effectively block TREM-2 signaling in primary PMN ^52^, recapitulated the increased IL-1β release compared to vehicle (DMSO) control-treated PMN (**Fig. 1E**). Next, we quantified the amount of ASC specks formation and found an increased number was present in XLA or BTK-inhibited PMN compared to HD PMN, both at basal conditions (**Fig. 1F, G**) and after LPS+nigericin stimulation (**Fig. 1H-I**). Switching to a genetic murine system and primary cells from *Btk*-deficient (KO) ^45^ or *Btk Xid*-mutated (*Xid*) mice ^53^ (see methods) we were able to confirm the data from human PMN vs PBMC/monocytes, namely that *Btk* KO or *Btk Xid* BMDM responded with significantly reduced IL-1β release (**Fig. 2A, B**), whereas BM-PMN showed significantly elevated IL-1β (**Fig. 2C, D**). Interestingly, LDH release was not affected (**Fig. S1I**). That IL-1β release was fully NLRP3-dependent in both murine cell populations was verified using the NLRP3 inhibitor MCC950 ^54^ (**Fig. 2A-D**) or *Nlrp3* KO BM-PMN and BMDM (**Fig. S1J, K**). This shows that in both human and mouse immune cells, BTK has a cell-type dependent rheostat function, augmenting nigericin-dependent IL-1β release in monocytes/macrophages but restricting it in neutrophils.

**Figure 1.**
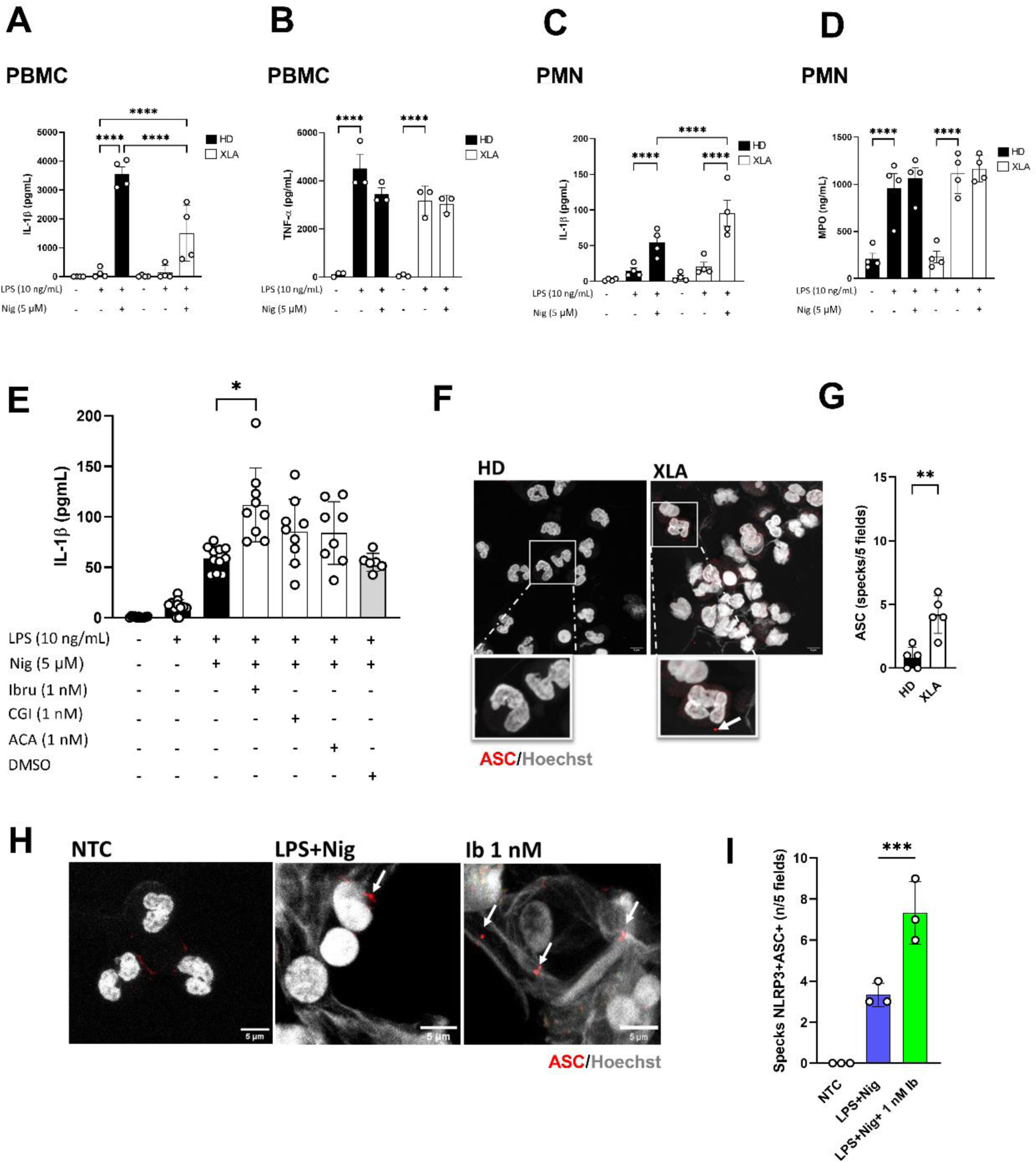
BTK ablation increases IL-1β release from primary human neutrophils but reduces IL-1β release from matched PBMCs. (**A**) IL-1β or (**B**) TNF-α release measured by triplicate ELISA in supernatants of stimulated PBMCs (A, B) isolated from HD and XLA (n = 4/4; each dot represents one donor); (**C**) IL-1β or (**D**) MPO release measured by triplicate ELISA in supernatants of stimulated peripheral blood PMNs isolated from HD and XLA (n = 4/4; each dot represents one donor); (**E**) IL-1β release in supernatants of peripheral blood PMN isolated from HD (n = 11); (**F**) ASC speck formation evaluated by confocal microscopy (grey: Hoechst staining for DNA, red: ASC staining, 63X magnification, arrows denote single specks) scale bar 1 µm; (**G**) Quantification ASC specks in 5 randomly chosen fields (one independent experiment); (**H**) ASC specks formation evaluated by confocal microscopy (grey Hoechst staining for DNA, red ASC staining, 63X magnification, arrows denote single specks) scale bar 5 µm; (**I**) Quantification of ASC specks in 3 randomly chosen fields (one independent experiment). (Each dot represents the average of technical triplicates per donor or biological replicate). A-D: Paired Two-way ANOVA test; E: Friedman test with multiple comparison correction; I: Paired one-way ANOVA with multiple comparison correction; G: Unpaired t-test, *p<0.05; **p<0.01; ***p<0.001; ****p<0.0001, HD: Healthy donors.

**Figure 2.**
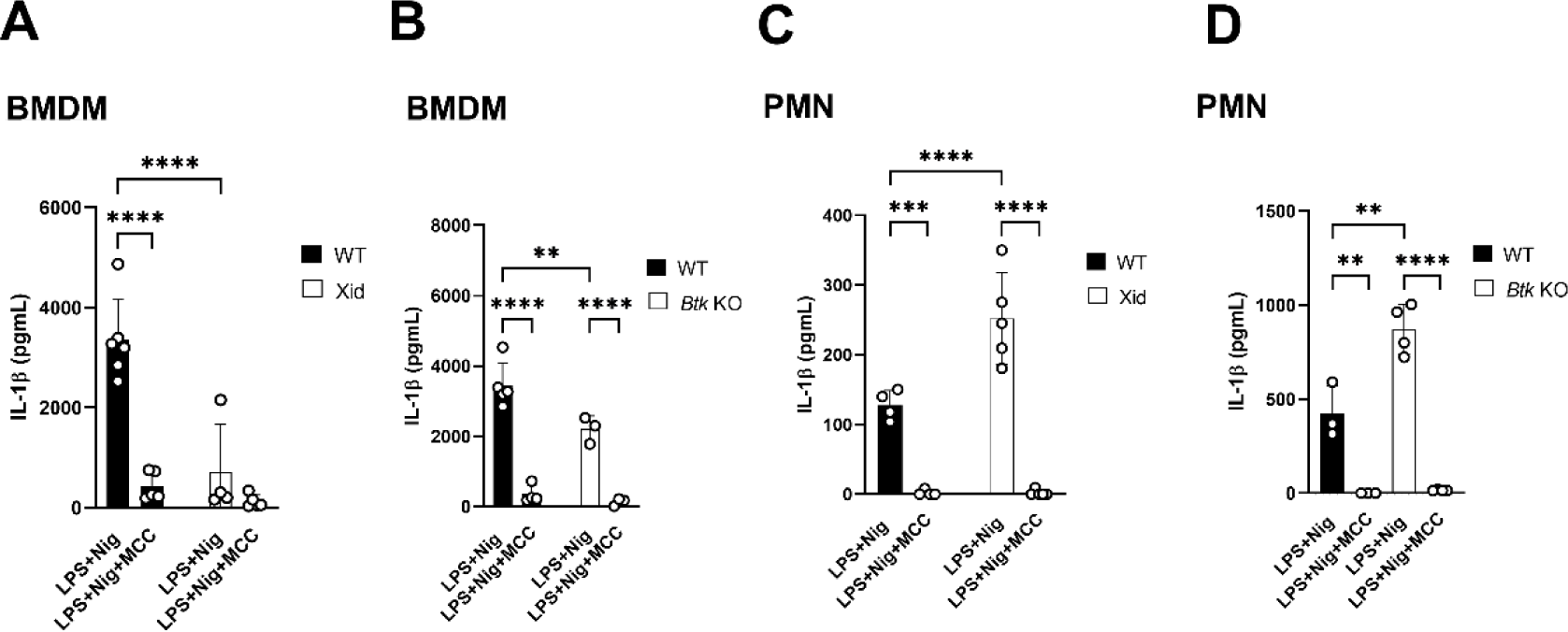
BTK ablation increases IL-1β release from primary bone-marrow derived neutrophils but reduces IL-1β release from matched BMDM. IL-1β release measured by ELISA in supernatants of stimulated BMDM from (**A**) *Xid* (n = 5), (**B**) *Btk* KO (n = 4), and in supernatant of BM-PMNs isolated from (**C**) *Xid* (n = 5), (**D**) *Btk* KO (n = 4). (**A-D**) C57BL6 WT (n = 9) mice (each dot represents one biological replicate). Paired One-way ANOVA test, *p<0.05; **p<0.01; ***p<0.001; ****p<0.0001.

### BTK restricts NLRP3-dependent NET release in primary PMN

The release of NETs, a pivotal role of PMN, was recently found to depend on NLRP3 ^4^. We therefore tested whether NET release might also be affected by the rheostat role of BTK by comparing NET release in HD vs XLA primary PMN or untreated with BTK inhibitor-treated primary PMN. Quantification of NETs by microscopy and MPO-dsDNA aggregate PicoGreen assay ^47^ showed that XLA PMN produced significantly more citrullinated H3 (citH3)-positive NETs (**Fig. 3A, B**) and MPO-dsDNA aggregates (**Fig. 3C**). Of note, NET release was entirely NLRP3-dependent as confirmed using the NLRP3 inhibitor MCC950, see **Fig. S2A, B**. Consistent with a higher responsiveness, we observed spontaneous NET release in XLA but not HD PMN in the absence of stimulation (**Fig. 3D, E**). Although they are generally less responsive than human primary PMN with regard to NET release, like HD human PMN, WT murine primary BM-PMN released significantly less NETs compared to *Btk Xid* and especially *Btk* KO BM-PMN (**Fig. 3F**, quantified in **3G**). *Nlrp3* KO PMN used as controls were nonresponsive as expected ^4^ and did not form NETs in response to LPS+nigericin. Thus, the negative regulatory role of BTK on NLRP3 activity in neutrophils extends to NET release.

**Figure 3.**
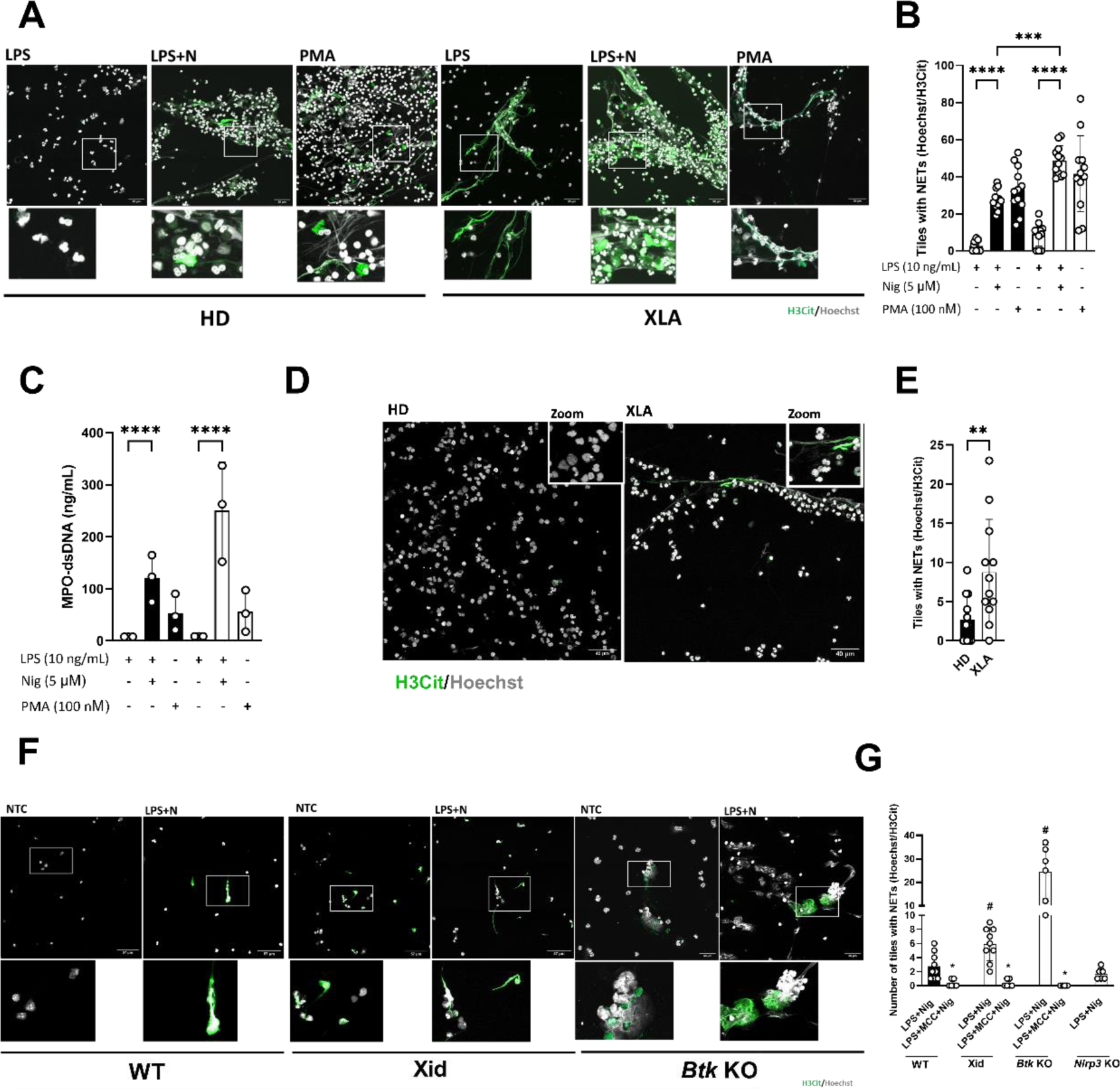
NET formation is dependent on NLRP3 inflammasome activation and increased in human and murine neutrophils when Btk is non-functional. (**A**) Representative NET formation measured by immunofluorescence imaging from healthy donors and XLA patients (grey: Hoechst representing DNA, green representing citrullinated histone H3); (**B**) NET quantification according to the number of tiles with NETs (Hoechst+H3Cit+), n = 3 HD and 3 XLA (each dot represents a random field from 3 independent experiments); (**C**) NET quantification (MPO-dsDNA) in neutrophils supernatant by Picogreen assay (each dot represents one independent experiment/donor); (**D**) Representative spontaneous NET formation measured by immunofluorescence imaging of PMNs from healthy donor and XLA patient (grey: Hoechst representing DNA, green representing citrullinated histone H3), scale bar 40 µm; (**E**) NET quantification according to the number of tiles with NETs (Hoechst+H3Cit+), n = 3 HD and 3 XLA (each dot represents a random field from 3 independent experiments/donors); (**F**) Representative NET formation measured by immunofluorescence imaging from *Xid*, *Btk* KO and WT mice (grey: Hoechst representing DNA, green representing citrullinated histone H3), scale bar 40 µm; (**G**) NET quantification according to the number of tiles with NETs (Hoechst+H3Cit+; each dot represents one biological replicate). B, C, G: Paired Two-way ANOVA test; E: Mann-Whitney; *p<0.05; **p<0.01; ***p<0.001; ****p<0.0001, HD: Healthy donors.

### BTK-NLRP3 regulated NET release involves canonical caspase-1

As certain details of PMN NLRP3 activation are currently controversial ^30, 31, 35^, we assessed whether the canonical NLRP3 executioner enzyme, caspase-1 ^55^, acted downstream of BTK/NLRP3. Of note, caspase-1 was activated by LPS+nigericin stimulation in primary PMN (**Fig. 4A, B**), like in monocytes/macrophages ^14, 19^. Unlike BTK, however, blockade of caspase-1 using the inhibitor YVAD ^56^ abrogated IL-1β release from both primary HD and XLA PMN (**Fig. 4C**) or WT vs *Btk* KO or *Btk Xid* BM-PMN (**Fig. 4D, E**). In human primary PMN release of MPO, a control readout, was not affected by caspase-1 blockade, as expected (**Fig. 4F**). Of note, NLRP3 inhibition with MCC950 also abrogated IL-1β release in all groups (**Fig. 4C-E**). Thus, neutrophil inflammasome employs canonical caspase-1 activity like in monocytes/macrophages despite the disparate role of BTK in both cell types.

**Figure 4.**
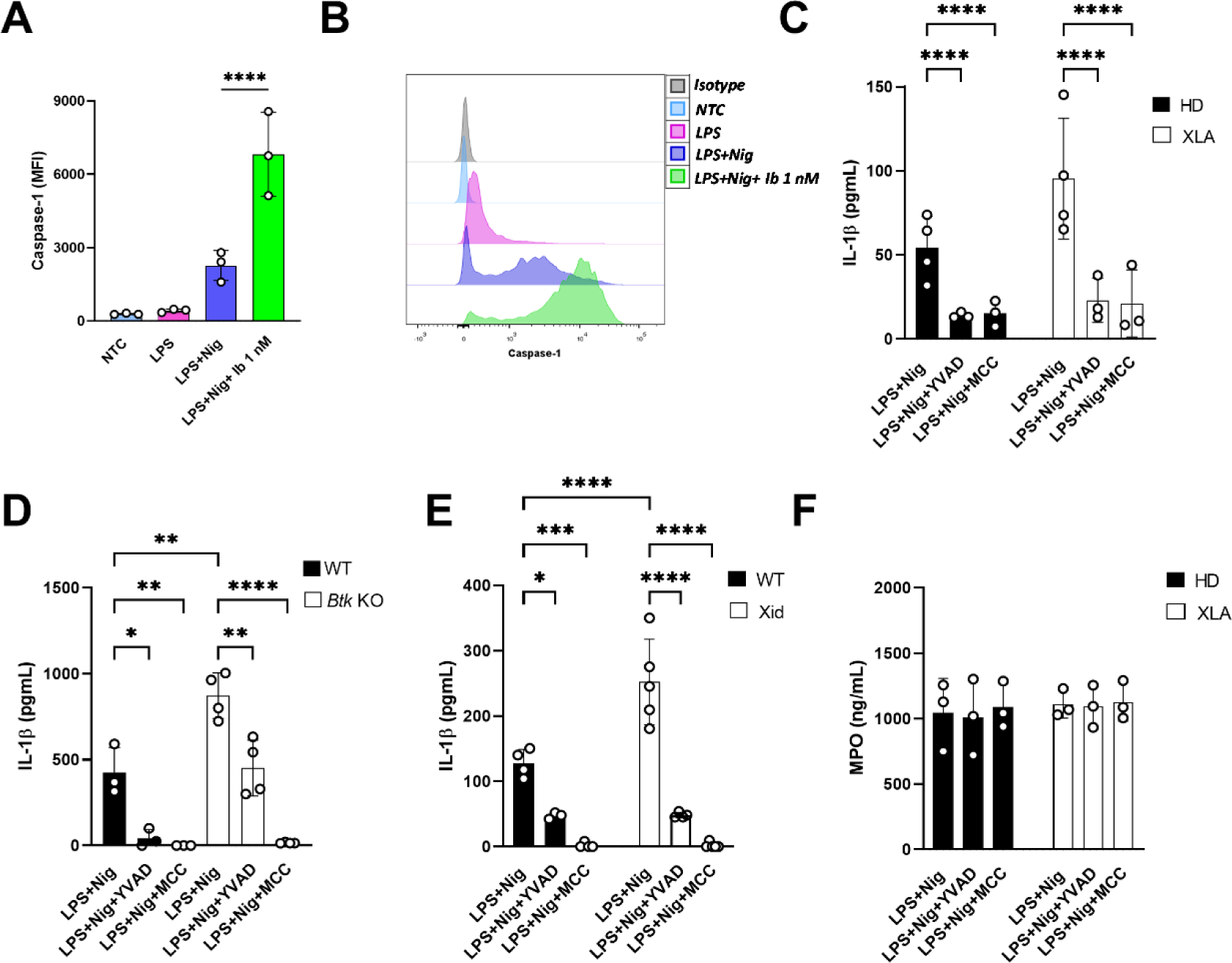
Caspase-1 role in BTK-dependent NLRP3 activation. (**A**) Median fluorescence intensity (MFI) and (**B**) representative overlayed histograms of caspase-1 activity in neutrophils stained with FAM-FLICA followed by flow cytometric analysis (each dot represents one independent experiment); IL-1β release measured by ELISA in the supernatant of (**C**) peripheral blood PMNs isolated from HD and XLA (n = 4/4; each dot represents one independent experiment); IL-1β release measured by ELISA in the supernatant of (**D**) BM-PMNs isolated from *Btk* KO (n = 4) and (**E**) *Xid* (n = 5) mice (each dot represents one biological replicate); (**F**) MPO release measured by ELISA in the supernatant of blood PMNs isolated from HD and XLA (n = 3/3; each dot represents one independent experiment). Paired Two-way ANOVA test, LPS+Nig versus inhibitors, *p<0.05; **p<0.01; ***p<0.001; ****p<0.0001, HD: Healthy donors. Ib = ibrutinib.

### MMP-9 is concomitantly an inflammasome-dependent alarmin and NLRP3 regulator

Given the recently emerged links between the inflammasome and azurophilic granule release (Johnson et al., 2017, ^57^), and more direct links between the tertiary granule cargo, MMP-9, and NLRP3 ^35, 38-40^, we sought to explore the dependence of this important matrix modifier on BTK regulation. Comparing MMP-9 release in primary HD vs XLA PMN we found significantly increased MMP-9 release for XLA PMN (**Fig. 5A**). In fact, MMP-9 levels correlated significantly with both IL-1β and NET release (**Fig. 5B, C**) with XLA PMN generally being more productive. Interestingly, carriers of the *NLRP3* gain-of-function variant rs10754558 variant ^49^ showed increased MMP-9 release (**Fig. 5D**). The dependence on BTK was also confirmed by ibrutinib inhibition in primary HD PMN (**Fig. 5E**). We also noted higher intracellular expression of MMP-9 which was boosted upon BTK inhibition (**Fig. 5F, G**). Since IL-1β had been suggested to be an MMP-9 substrate ^41, 42^, we speculated that MMP-9 might not only be an NLRP3-dependent secreted factor but also contribute proteolytically to NLRP3 inflammasome activity. To address this, the selective MMP-9 inhibitor, JNJ ^58^, was used in primary human PMN. Interestingly, JNJ pre-treatment strongly blocked MMP-9 but also IL-1β release upon LPS+nigericin treatment), without affecting the NLRP3-independent release of MPO (**Fig. 5H-I**). Moreover, NET release (assessed by both microscopy and MPO-dsRNA analysis) was strongly impaired (**Fig. 5 J-L**). This data suggests that in human neutrophils, not only caspase-1, but also MMP-9 is involved in IL-1β processing and NET formation downstream of NLRP3. To reinforce this theory, we also investigated the caspase-1 activity upon MMP-9 inhibition, and we did not observe a significant effect on caspase-1 activation, suggesting that indeed MMP-9 role could be downstream NLRP3/caspase-1 (**Fig. S3**). Thus, MMP-9 appears to be both alarmin and executioner of NLRP3 activity in neutrophils.

**Figure 5.**
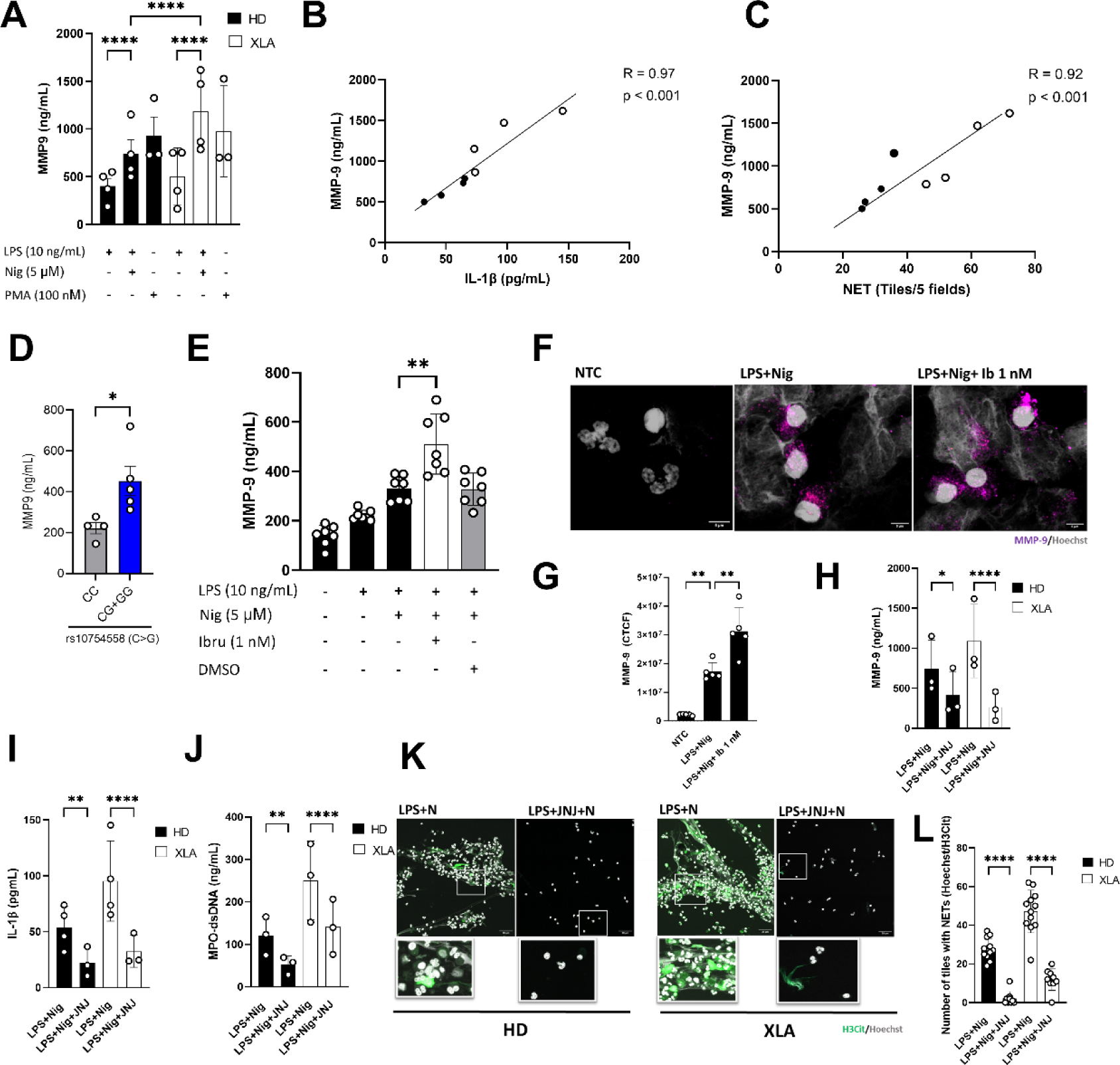
MMP-9 release is increased upon NLRP3 inflammasome activation. (**A**) MMP-9 release measured by ELISA in the supernatants of peripheral blood PMNs isolated from HD and XLA (n = 4/4; each dot represents one independent experiment); (**B**) Spearman correlation analysis of MMP-9 and IL-β release or (**C**) MMP-9 and NET release from neutrophils isolate from HD and XLA (n = 4/4); (**D**) MMP-9 levels in HD (n = 10) grouped according to *NLRP3* SNV genotype (rs10754558, C>G); (**E**) MMP-9 release in the supernatants of peripheral blood primary neutrophils isolated from HD (n = 11); (**F**) Representative MMP-9 expression measured by immunofluorescence imaging from healthy donor treated or not with Ibrutinib 1 nM (grey: Hoechst representing DNA, magenta representing MMP-9 staining, scale bar = 5 µm); (**G**) Corrected total cell fluorescence (CTCF) for MMP-9 in 5 random fields; (**H**) MMP-9 or (**I**) IL-1β release measured by ELISA in the supernatant of blood PMNs isolated from HD and XLA (n = 4/4); (**J**) NET quantification (MPO-dsDNA) in neutrophils supernatant by Picogreen assay (each dot represent one independent experiment); (**K**) Representative NET formation measured by immunofluorescence imaging from healthy donor and XLA patient (grey: Hoechst representing DNA, green representing citrullinated histone H3, scale bar = 38 µm); (**L**) NET quantification according to the number of tiles with NETs (Hoescht+H3Cit+). A, H, I, J, L: Two-way ANOVA test; B, C: Spearman correlation analysis; D: Mann-Whitney test; G: Paired one-way ANOVA with multiple comparison; *p<0.05; **p<0.01; ***p<0.001; ****p<0.0001, HD: Healthy donors.

## Discussion

The results obtained in this study raise the intriguing idea that BTK differentially regulates NLRP3 inflammasome activation depending on the cell type: In PBMCs or BMDMs, it was previously described that BTK interacted with NLRP3 and phosphorylated several conserved tyrosine residues, which influenced NLRP3 cellular localization, oligomerization and activation ^19^. Thus, in both human monocytes and murine macrophages BTK was described as a positive regulator of NLRP3 inflammasome activation ^14, 19, 23^. This is also in line with human data ^59^ and mouse in vivo models ^25, 60, 61^.

We were able to replicate these observations in human PBMCs and murine BMDM; however, to our surprise, in primary neutrophils this regulation occurred in an opposite direction. For the first time we demonstrate here that BTK might have a negative regulatory role on NLRP3 activation in both, human and murine neutrophils. Even though low LPS (10 ng/ml) was used here to prime neutrophils, this unexpected property is somewhat reminiscent of Mao et al ^27^, who observed that BTK could switch from a positive regulator at low LPS concentrations to a negative regulator at high LPS concentrations in macrophages/monocytes (for comment see ^62^). Given the high reactivity of PMN, one could speculate that in PMN BTK is therefore already “pre-configured” to only negatively regulate the NLRP3 inflammasome, consistent with an increased basal IL-1β and NET release. The molecular basis for this is unclear. Mao et al proposed BTK-mediated phosphorylation (and hence de-activation of) of the phosphatase PP2A might negatively regulate NLRP3 in macrophages, although some of this analysis was carried out in HEK293T cells ^27^. In this scenario, PP2A promotes NLRP3 activity by removing an inhibitory Ser 5 phospho-site in NLRP3 ^51^. Consequently, BTK would inhibit NLRP3 by preventing PP2A-mediated dephosphorylation at Ser 5. Therefore, BTK loss or inhibition would increase PP2A activity, rending NLRP3 more reactive. If the molecular basis for this proposed inhibitory role of BTK in macrophages applied to neutrophils, then a tonic BTK activity might keep PP2A (and hence NLRP3) in check by maintaining a pool of inactive p-PP2A, a mechanism to explore in the future. Alternatively, NLRP3 hyperactivation might relate to increased reactive oxygen species (ROS) production observed in *Btk*-deficient neutrophils due to the lack of negative regulation imposed on NADPH oxidase ^63^. However, the effect of ROS on NLRP3 activation has recently been challenged ^64^. Unfortunately, due to limitations related to the access and rarity of XLA samples and the poor availability of PP2A phospho-antibodies ^51^, testing these scenarios experimentally was outside the scope of this present study. Nevertheless, we have characterized the effect of BTK in NLRP3 activation in neutrophils in different primary cell models (mouse vs human, genetic vs inhibitor treatment), providing new insights into neutrophil inflammasome biology.

Moreover, our results highlight how the effect of BTK inhibitors may be cell-context-dependent: whereas macrophages NLRP3 activity is blocked by BTK inhibitors, neutrophil NLRP3 is boosted. This is highly relevant, considering that some BTK inhibitors are FDA approved (i.e.: ibrutinib and acalabrutinib) or are currently used in several clinical trials and treatments for inflammatory and malignant diseases ^65^. Therefore, understanding the effect of these drugs on a cellular level might help to guide different clinical approaches in different inflammasome-related diseases, depending e.g. on the dominance of macrophage vs neutrophil involvement. Our study suggests that BTK inhibition may be double-edged: dampening monocyte-mediated inflammation as evidenced ex vivo in ibrutinib-treated patient PBMC (Liu et al., 2017 ^14^), whilst augmenting PMN-mediated inflammation. As the production of at least IL-1β from monocytes far exceeds IL-1β released from PMN, the net result of BTK inhibition in patients, at least for IL-1β, may still be a restriction of inflammation. However, how IL-1β inhibition balances with the strong positive effect of BTK inhibition on NET release is unclear. Of note, NETs contain inflammasome-activating DAMPs such as naRNA-LL37 complexes that drive further NLRP3-dependent NETosis ^66^ so that increased NET formation due to BTK inhibition may fuel inflammation overall, at least transiently. On the other hand, in XLA patients, increased neutrophil NLRP3 activity and increased NET formation may compensate for the described ineffectiveness of NLRP3 activity in macrophages and the lack of B cells and antibodies ^14^. Further clinical studies in both XLA patients and patients receiving BTK inhibitors may have to carefully investigate this.

Interestingly, other neutrophil proteases, e.g. elastase, have been involved in IL-1β processing ^67^. In addition, we provide evidence that in neutrophils, the proteolytic activity of caspase-1 may be complemented by other enzymes like MMP-9 for IL-1β maturation. This data, at least *in vitro,* is consistent with previous findings that showed that MMP-9 release is dependent of NLRP3 activation, as well as its association with a dysregulated NLRP3 inflammasome in inflammatory conditions such as severe COVID-19 ^35^, models for aortic aneurism ^39^, renal inflammation ^68^, retinopathy ^69^ and intracranial aneurism ^40^. We can only speculate how MMP-9 is activated upon NLRP3 inflammasome activation. We hypothesize that a protease like caspase-1 could act as an intermediate player between NLRP3 activation and MMP-9 activity. Ren et al., 2020 suggested that caspase-1 could directly activate MMP-9 by cleaving its N-terminal inhibitory domain leading to its activation. Indeed, when we specifically inhibited caspase-1 (YVAD treatment) we observed a significant reduction of MMP-9 expression and release, suggesting that MMP-9 could be downstream caspase-1 activation.

Collectively, our study highlights BTK and MMP-9 as novel regulators of the neutrophil NLRP3 inflammasome. This will have implications especially for the clinical use of BTK inhibitors. Further studies are required to better understand the dynamics of BTK regulation, NLRP3 activation and the downstream caspase-1/MMP-9/IL-1β axis/NET axis in neutrophils, not least as targeting these pathways has important ramifications for inflammatory disease.

## Abbreviations

AF: Alexa Fluor
DAMP: damage-associated molecular pattern
FACS: Fluorescence Activated Cell Sorting
IFN: Interferon
IL: Interleukin
KO: knockout
MMP: matrix metalloproteinase
MPO: Myeloperoxidase
NET: Neutrophil extracellular trap
NLRP3: NOD-like receptor family, pyrin domain containing 3
PAD4: Peptidyl arginine deiminase 4
PBMC: Peripheral Blood Mononuclear Cell
PMN: Polymorphonuclear leukocytes
TLR: Toll-like receptor
TNF: Tumor necrosis factor

## Acknowledgements

We thank Libera Lo Presit for editorial support, Maria Mateo Tortola for support regarding microscopy and Markus W. Löffler for biobank support and acquisition of blood samples. The study was supported by the grants from the internal support program of the Medical Faculty, University of Tübingen, and the DFG (German Research Foundation) research grant We-4195/18-1 (to A. N. R. W), CRC TR156 “The skin as an immune sensor and effector organ – Orchestrating local and systemic immunity” (project B05 to FB and ANRW), the Clusters of excellence “iFIT-Image Guided and Functionally Instructed Tumor Therapies” (EXC-2180, to A.N.R.W) and “CMFI-Controlling Microbes to Fight Infections” (EXC-2124, to A. N. R. W.). VNCL was supported by grant 2021/13049-4 and 2020/15323-3 of the São Paulo Research Foundation (FAPESP).

## Author contributions

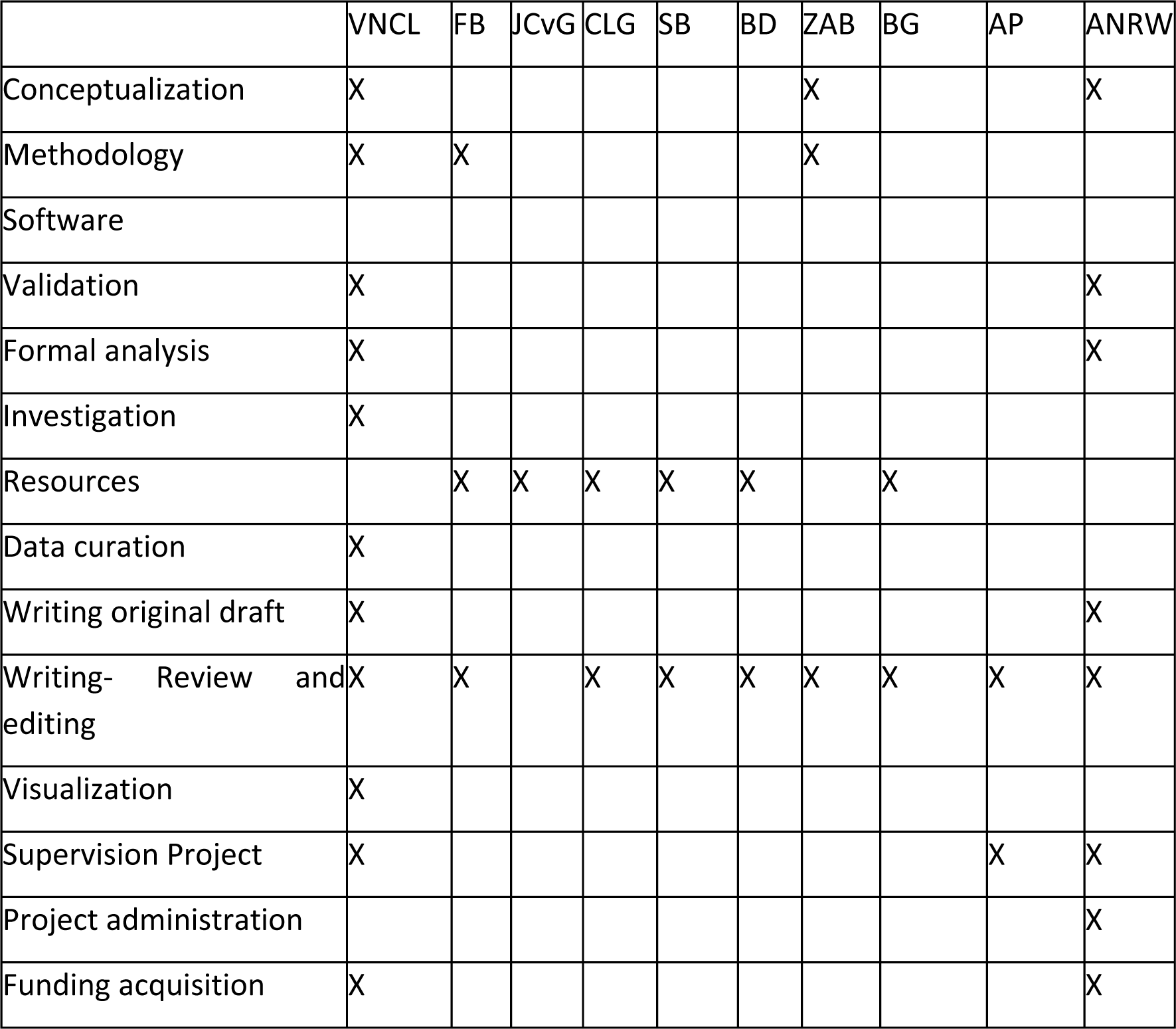

## SUPPLEMENTARY FILES

**Supp Fig 1.**
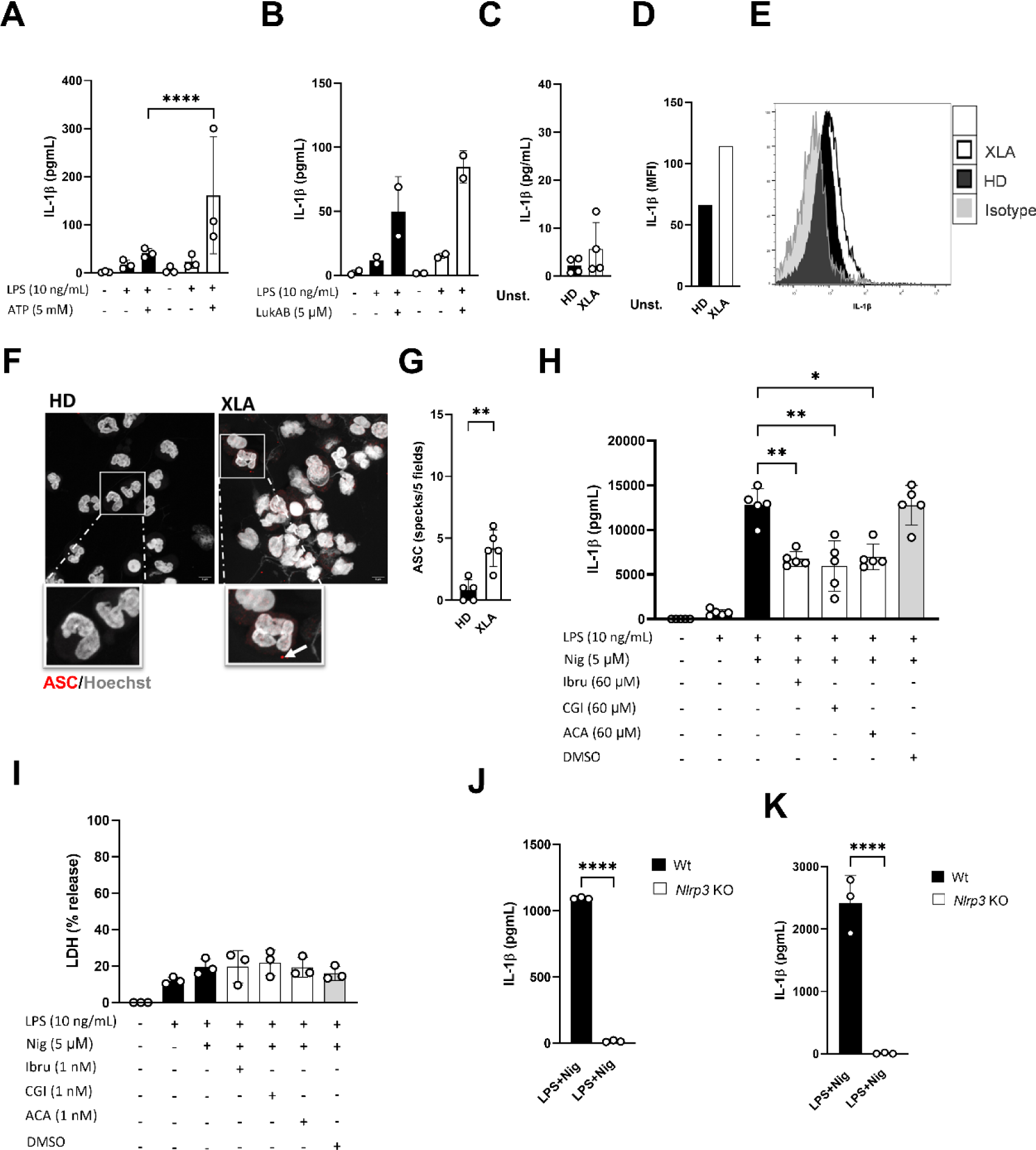
IL-1β release measured by ELISA in supernatant of peripheral blood PMNs isolated from HD and XLA (n = 3/3) and stimulated with (**A**) ATP or (**B**) LukAB, each dot represents one independent experiment); (**C**) Basal IL-1β measured by ELISA in supernatant of peripheral blood PMNs isolated from HD (n = 4) and XLA (n = 4); (**D**) Intracellular IL-1β (MFI) quantified by FACS in peripheral blood PMNs (CD66b+CD11b+CD15+) isolated from HD (n = 1) and XLA (n = 1), (**E**) Representative histogram of intracellular IL-1β quantified by FACS in peripheral blood PMNs; (**F**) ASC “specks” formation evaluated by confocal microscopy (grey Hoechst staining for DNA, red ASC staining, 63X magnification, arrows denote single specks), scale bar 5 µm; (**G**) Quantification ASC specks in 5 random fields (one independent experiment); (**H**) IL-1β and (**I**) LDH (%) release in supernatant of peripheral blood mononuclear cells (PBMC), n = 5 HD; IL-1β release measured by ELISA in supernatant of (**J**) peripheral blood PMNs and (**K**) BMDM isolated from *Nlrp3* KO mice (n = 3/3). A, B: Two-way ANOVA test; C, G: Mann-Whitney test; J, K: t-test; H, I: Paired one-way ANOVA with multiple comparison; *p<0.05; **p<0.01; ***p<0.001; ****p<0.0001, HD: Healthy donors.

**Supp Fig 2.**
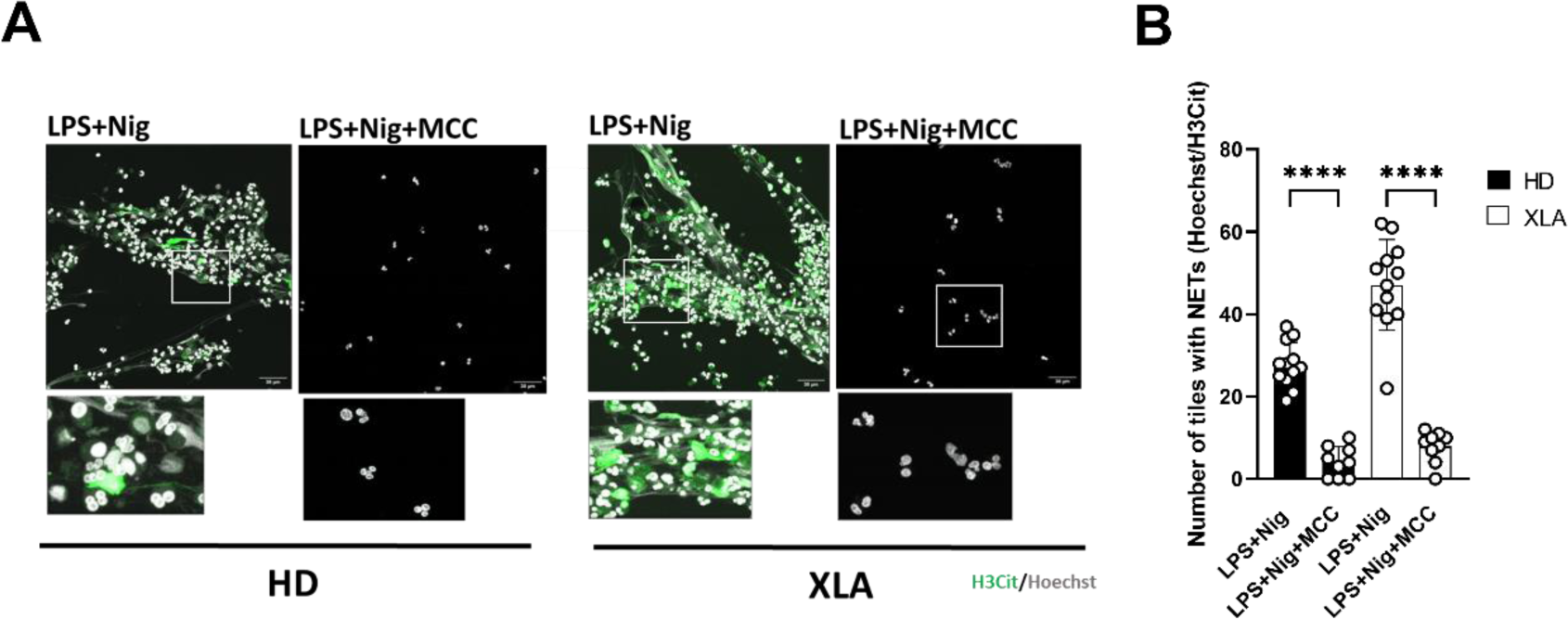
(**A**) Representative NET formation measured by immunofluorescence imaging from healthy donor and XLA patient (grey: Hoechst representing DNA, green representing citrullinated histone H3, scale bar = 40 µm); (**B**) NET quantification according to the number of tiles with NETs (Hoechst+H3Cit+). Two-way ANOVA test; *p<0.05; **p<0.01; ***p<0.001; ****p<0.0001, HD: Healthy donors.

**Supp Fig 3.**
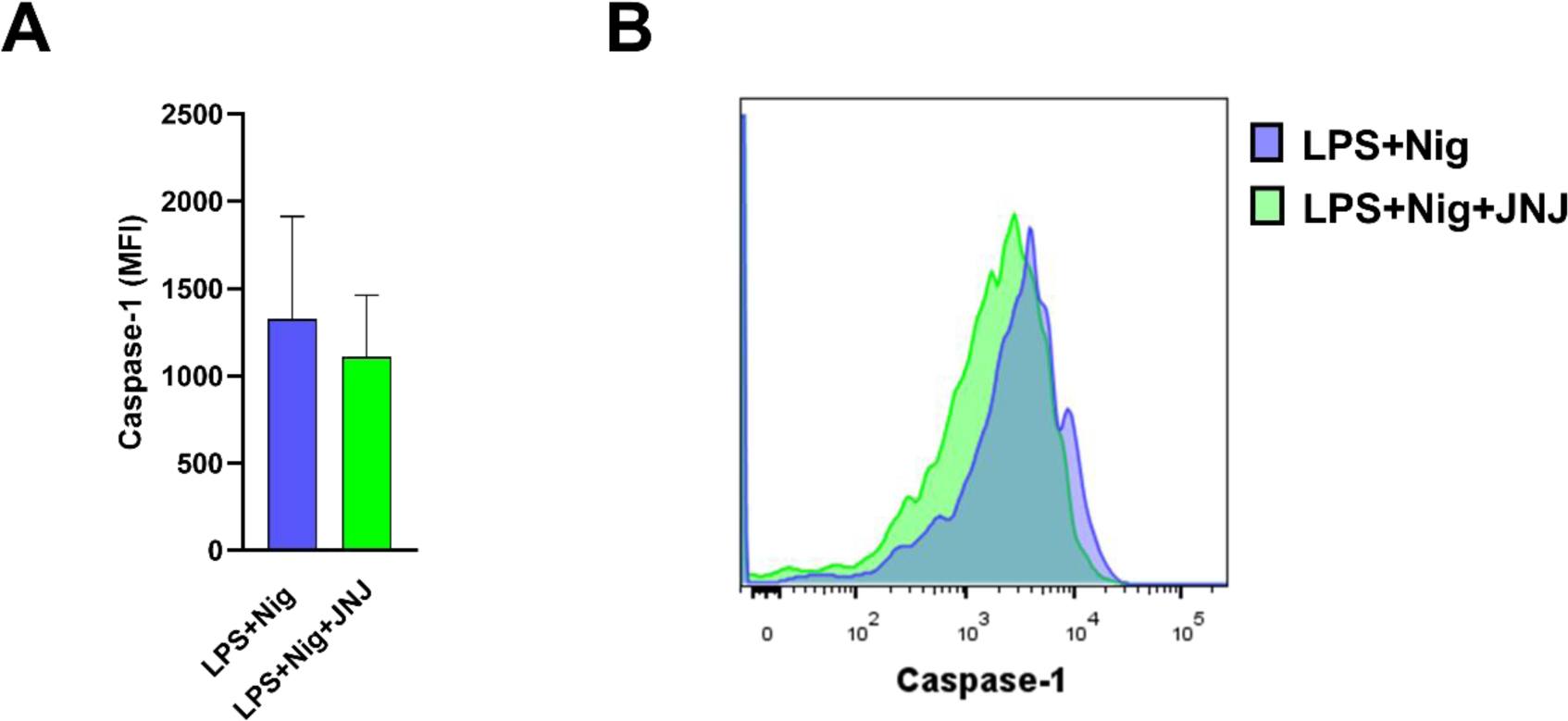
(**A**) Median fluorescence intensity (MFI) and (**B**) Representative overlayed histograms of caspase-1 activity in neutrophils stained with FAM-Flica and flow cytometric analysis (each dot represents one independent experiment); Paired “t” test, p>0.05.

**Table S1:** Materials and Kits used.

| Name | Company | Cat |
| --- | --- | --- |
| EDTA pH 8 | Sigma | E9884 |
| Ampuwa | Fresenius Kabi | 1833 |
| RPMI | Sigma-Aldrich | R8758 |
| OptiMEM | Gibco, Thermo | 11058021 |
| FCS | TH-Geyer | 11682258 |
| BSA | BIOMOL | 1.400.100 |
| Ficoll | Sigma-Aldrich | 10771 |
| DEPC (Diethyldicarbonate) | Roth | K028.2 |
| Saponin | Applichem | A4518.0100 |
| Fixation Buffer containing 4% Paraformaldehyde | Biolegend | 420801 |
| PBS (Phosphate buffered saline) | Thermo Fisher | 14190-169 |
| ProLong Diamond Antifade Mountant | Thermo Fisher | P36965 |
| Chicken serum | Vector Laboratories | S-3000 |
| Hoechst3334 | Thermo Scientific | 62249 |
| DuoSet® ELISA human MMP-9 | R&D Systems | DY911-05 |
| DuoSet® ELISA human MPO | R&D Systems | DY3174 |
| ELISA MAX™ Deluxe Set Human IL-1β | Biolegend | 437004 |
| ELISA MAX™ Deluxe Set Human IL-8 | Biolegend | 431504 |
| TNF alpha Human Uncoated ELISA Kit | ThermoFisher | 88-7346-22 |
| ELISA MAX™ Deluxe Set Human IL-6 | Biolegend | 430504 |
| R&D Systems™ Mouse IL-1 beta/IL-1F2 DuoSet ELISA | R&D Systems | DY40105 |
| ELISA MAX™ Deluxe Set Mouse TNF-α | Biolegend | 430904 |
| Cytotoxicity Detection Kit (LDH) | Roche | 11644793001 |
| Quant-iT™ PicoGreen™ | Thermo Fisher | P11495 |
| Neutrophil Isolation Kit, mouse | Miltenyi Biotec | 130-097-658 |
| LPS, E. coli 0111:B4 | Invivogen | tlrl-3pelps |
| Nigericin | Invivogen | tltl-Nig |
| Adenosine 5'-triphosphate disodium salt hydrate, ATP | Sigma | A2383 |
| JNJ0966 | MedChemExpress | HY-103482 |
| MCC950 | Cayman | 17510 |
| YVAD | Adipogen | Inh-Yvad |
| R406 | Selleckchem | S1533 |
| Disulfiram | MedChem | HY-B0240 |
| Ibrutinib | Selleckchem | PCI-32765 |
| CGI | Selleckchem | S7051 |
| Acalabrutinib | Selleckchem | S8116 |

**Table S2:** Buffers and Recipes.

| Buffer/Medium | Recipe |
| --- | --- |
| Erythrocyte Lysis Buffer 10x | 1.54 M NH<sub>4</sub>Cl, 100 mM KHCO<sub>3</sub>, 1 mM EDTA pH=8, Dissolved in Ampuwa sterile water, pH adjusted = 7.3, sterile filtered (0.22 µm). |
| FACS Buffer | PBS, 2% FCS heat inactivated, 1 mM EDTA pH=8, Sodium Azide 0.1% |
| Neutrophil culture medium | RPMI, 5% FCS heat inactivated sterile filtered (0.22 µm), 1% L-Glutamin |
| Blocking Buffer IF | PBS, 10% Chicken serum, 0.1% Saponin |
| Blocking solution-Picogreen | PBS, 2% BSA |
| Washing solution-Picogreen | PBS, Tween 20 (pH 7.4) |
| TE Buffer 1x-Picogreen | 20x TE buffer provided by kit 1:20 in DNase free H<sub>2</sub>O |
| IF wash solution | PBS, 0.05% Saponin |
| MACS Buffer | PBS, 0.5% BSA, 2 mM EDTA |
| BMDM Culture medium | RPMI, 10% FCS, 1% L-glutamine, 1% Penicillin-Streptomycin |
| BM Medium | RPMI, 10% FCS |

**Table S3:** Antibodies.

| Name | Company | Cat. | Dilution |
| --- | --- | --- | --- |
| Anti-myeloperoxidase | ThermoFisher | PA516672 | 1:500 |
| Anti-H3Cit | Abcam | AB5103 | 1:200 |
| Anti-ASC | Santa Cruz | sc271054 | 1:200 |
| Anti-MMP-9 | Invitrogen | MA5-15886 | 1:200 |
| Anti-Btk | Cell Signalling | #8547 | 1:200 |
| Alexa 594-labeled chicken anti-mouse | ThermoFisher | A-21201 | 1:500 |
| Alexa 647-labeled chicken anti-mouse | ThermoFisher | A21463 | 1:500 |
| Alexa 594-labeled chicken anti-rabbit | ThermoFisher | A21442 | 1:500 |
| Alexa 594-labeled chicken anti-goat | ThermoFisher | A21468 | 1:500 |
| Alexa 488-labeled goat anti-rabbit | ThermoFisher | A11034 | 1:500 |
| Pacific Blue-labeled anti-human CD14 (mouse) | Biolegend | 325616 | 1:50 |
| PE-labeled anti-human CD14 (mouse) | Biolegend | 367103 | 1:50 |
| FITC-labeled anti-human CD66b (mouse) | Biolegend | 305104 | 1:50 |
| APC-labeled anti-human CD11b (mouse) | Biolegend | 301310 | 1:50 |
| APC-Cy7-labeled anti-human CD11b (mouse) | BD | 557754 | 1:50 |
| PE-labeled anti-human CD15 (mouse) | Biolegend | 323006 | 1:50 |
| PE-Cy7-labeled anti-human CD15 (mouse) | Biolegend | 323030 | 1:50 |
| BV421-labeled anti-human CD62L (mouse) | Biolegend | 304828 | 1:50 |
| PE-labeled anti-human IL-1β (mouse) | Biolegend | 12-7018-41 | 1:20 |
| APC-labeled anti-mouse LY6C/G (rat) | Biolegend | 108412 | 1:50 |
| FITC-labeled anti-mouse CD11b (rat) | BD | 553310 | 1:50 |
| FITC-labeled IgG1 isotype (mouse) | Miltenyi | 130-113-761 | 1:50 |
| PE-labeled IgG1 isotype (mouse) | ImmunoTools | 21275514 | 1:50 |
| APC-labeled IgG1 isotype (mouse) | Miltenyi | 130-112-758 | 1:50 |
| FITC-labeled IgG2a isotype (mouse) | ImmunoTools | 21275523 | 1:50 |
| FITC-labeled IgM isotype (mouse) | Biolegend | 401606 | 1:50 |
| PECy-7-labeled IgG1 isotype (mouse) | Biolegend | 400126 | 1:50 |

